# Entropic regulation of dynamical metabolic processes

**DOI:** 10.1101/643601

**Authors:** Stephan O. Adler, Edda Klipp

## Abstract

Life depends on the input of energy, either directly provided by sunlight or in form of high-energy matter. The rules and conditions for the conversion of chemical or electromagnetic energy into living structure and all the processes related with life are governed by the laws of thermodynamics. Hence, to understand the potential and the limitations of cell growth and metabolism, it is unavoidable to take these laws into account. During the last years, systems biology has developed many mathematical models aiming to describe steady states and dynamic behavior of cellular processes in qualitative and quantitative terms. The validity of the model predictions depends strongly on whether the model formulation is in agreement with the laws of physics, chemistry, and, specifically, thermodynamics.

Here, we review basic principles of thermodynamics for equilibrium and non-equilibrium processes as well as for closed and open systems as far as they concern metabolic processes, especially in their dynamics. We illustrate the application of thermodynamic laws for some practical cases that are currently intensively studied in systems and computational biology. Specifically, we will discuss the concept of entropy production and energy dissipation for isolated and open systems and its interpretation for the feasibility of biological processes, especially metabolism. We demonstrate that steady states of metabolic systems cannot show energy dissipation, while in dynamical modes entropy of the system can be both increased or decreased, depending on the type of perturbation and the kinetics of the reaction system. These findings are very important for biotechnological processes where energy dissipation should be limited, but also for analysis of healthy and diseased cellular metabolism.

## Introduction

Processes of life are dynamical, at all levels from the evolution of species and the development of a single higher organism down to cellular organization and maintenance including the cell cycle. A plethora of mathematical models have been developed during the last decades explaining the observed behavior for cellular processes such as the cell cycle, signaling or metabolism as well as for processes in tissues, organs, organisms or between species.

Mathematical modeling of metabolism here takes a special role. Many observed phenomena could be explained by steady state models assuming constant concentrations of metabolites despite non-zero fluxes between these metabolites. This, together with the formulation of an objective function, is the basis for an approach known as flux balance analysis. Adding constraints to fluxes or other observables then leads to constraint-based modeling. This approach has been very fruitful for a number of important findings^1–4^. However, it must ignore temporal changes, which are often relevant. Non-equilibrium thermodynamics allows to tackle processes in time, to determine directionality, and to assess the energy spend on dynamical trajectories.

Below, we will first revisit the basic principles of phenomenological equilibrium thermodynamic descriptions. Then we will consider open systems and steady states as well as deviation from steady states. Applications comprise biotechnological processes such as chemostats and batch culture, but also metabolic changes between healthy cells and cancerous mutations.

## Basic Principles

### Systems, states, variables

We first have to define a number of notions. As you will see they are in part mutually dependent on each other. The first important term that we need is the *system*. A system is here every physical entity that can be distinguished from its environment. The *environment* is everything else that can interact with the system. The discrimination between system and environment can be spatial but does not need to be.

Practically useful examples for systems and their environment are (i) a chemical reaction and the solution, (ii) a microbial cell and its growth medium, or (iii) the mitochondrion and the cytoplasm. Depending on the type of exchange between system and surroundings, we discriminate between open and closed systems. Open systems exchange matter and energy, closed systems only exchange energy. Finally, for isolated systems the boundary permits neither energy nor matter exchange. If *U* is the internal energy and *M* the mass of the system, we can write:

Isolated system: d*U* = 0, d*M* = 0

Closed system: d*U* ≠ 0, d*M* = 0

Open system: d*U* ≠ 0, d*M* ≠ 0

Here, the “d” marks any perturbation that can happen in agreement with the general conditions.

Living systems are all open systems. They require input of energy and matter to sustain their structure, for growth and development.

Thermodynamic systems have *temperature T* as a property. Typically, they consist of many objects such as atoms, particles, molecules. If we are interested in describing the interaction of these particles, i.e. by considering their kinetic and interaction energies, we employ the microscopic treatment used in *statistical thermodynamics*. This will not be further discussed here. *Phenomenological thermodynamics* uses a macroscopic treatment: we consider measurable properties that belong to the system as a whole such as temperature *T*, pressure *p*, volume *V*, internal energy *U*. Further frequently used variables are concentrations *c*_*i*_, charges *z*_*i*_, or electrical fields. Variables required to describe the state of the system are *state variables*. The value of a state variable is only dependent on the state of the system, not on the path taken to arrive at this state. Since there are dependencies between variables belonging to a state of a system, there is a smallest set of (independent) state variables. The *state* is a snapshot of the thermodynamic system described by the state variables.

Four important laws form the basis of thermodynamic research, two are of practical importance in biology. The so-called *first law* states that the state variable internal energy *U* changes through uptake of heat *Q* or work *W*:

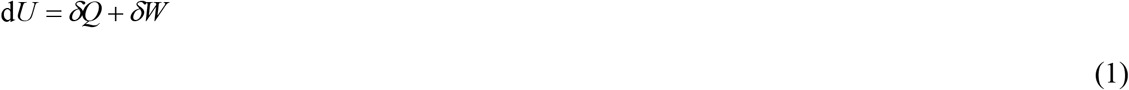

(we use *δ* instead of d to indicate that *Q* and *W* are not state variables; their values also depend on the path on which the state was reached). For many practically relevant cases, we can restrict the consideration of work to volume work replacing *δW* by -*p*d*V* (i.e. change of internal energy due to compression or dilatation).

The *second law* introduces the entropy *S* as a new state variable. If the system is in thermodynamic equilibrium with the environment, then we can calculate the changes of *S* during reversible processes in the following way:

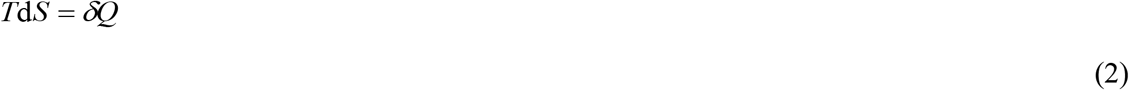

Combining equations (1) and (2) leads to *Gibbs’ fundamental law*

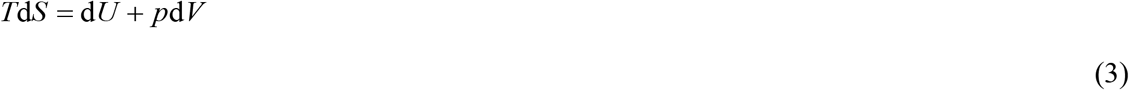

Thus, the variation of entropy is a measure for the exchange of energy with the environment in reversible processes.

Experience has taught that every isolated thermodynamic system evolves towards a state that it cannot leave spontaneously, the *equilibrium state*. During this irreversible process, entropy can only increase. The equilibrium state is characterized by a maximum of the entropy and a minimum of Gibbs free energy of the system. Due to their importance for states and dynamics of biochemical or biological systems, we will discuss properties of both quantities below.

### The Gibbs Free Energy

Among the thermodynamic potentials, Gibbs free enery (also known as free enthalpy) *G* is mostly suited to describing biochemical reactions systems. It is related to internal energy *U*, enthalpy *H* and entropy *S via* the relation *G* = *H* − *TS* = *U* + *pV* − *TS*. As thermodynamic potential, *G* is a function of pressure *p* and temperature *T*, i.e. *G* = *G*(*T*, *p*). In equilibrium, *G* assumes a minimum, if *p* and *T* are kept constant. For multi-component systems, it is also dependent on the mole numbers *n*_*i*_ of all components *i* (*i* = 1,…, *K*) and it holds the Gibbs-Duhem equation

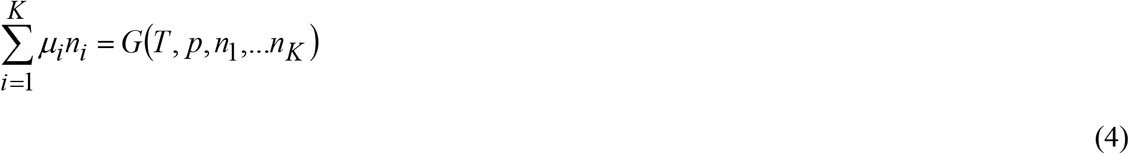

holds, which establishes a relation among the Gibbs enthalpy and the chemical potentials *μ*_*i*_ of the components. The other way around, the chemical potential expresses the change of *G* with changing mole numbers:

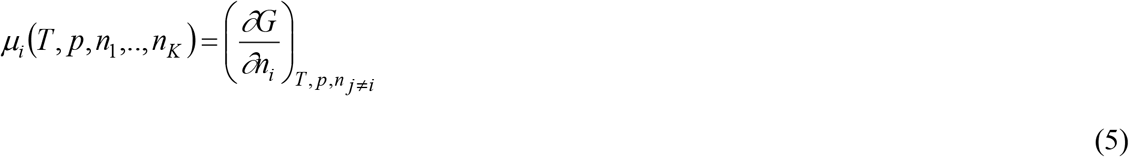

In the analysis of biochemical reaction networks, we are often interested in the changes of *G* (e.g. due to reactions or substance uptake/release) or in its differences. Here, it is important to distinguish between various differences in literature:

i. The difference of Gibbs enthalpies for two distinct system states, denoted here with Δ_*s*_*G*. In equilibrium thermodynamics, it is related to the respective differences of the other thermodynamic potentials, e.g. Δ_*s*_*G*= Δ_*s*_*H* − *T*Δ_*s*_*S* for constant temperature.
ii. The Gibbs enthalpy of formation, here denoted with Δ_*f*_*G*_*i*_ for components *i*. Since *G* is a state variable, it is independent of how the state was reached. For practical purposes, the Gibbs enthalpy of component *i* can be calculated from the Gibbs enthalpy of its constituents and the enthalpy required to create the considered component.
iii. The difference of Gibbs enthalpies of educt and product of a biochemical reaction, Δ_*r*_*G*. For reaction A ↔ B, it holds that Δ_*r*_*G*_A↔B_ = Δ _*f*_*G*_*A*_− Δ _*f*_*G*_*B*_. The reaction enthalpy is also related to the equilibrium constant *K_eq_* of that reaction, i.e. 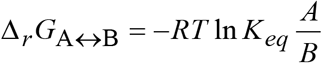 with *A* and *B* being the concentrations of A and B, respectively. For most practical cases, the reaction enthalpy can also be related to the affinity of a reaction, *A*_*r*_, by 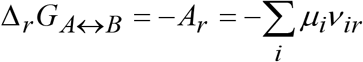 where the *v*_*ir*_ are the stoichiometric coefficients of substance *i* in reaction *r* and, hence, the sum extends over all educts and products of the considered reaction.

Up to now, we have only considered concepts of *equilibrium thermodynamics*, which deals with the description of state variables and their dependencies in equilibrium. It also studies the differences of state variables when comparing two different equilibrium states. It essentially ignores the *process* through which the system evolves from one state to the other. Although most of the laws and principles discussed above have originally been derived for isolated systems, they also have implications for or are practically applicable to open systems, as we will demonstrate. However, it is important to keep track of uptake, conversion and release of the different quantities such as energy and matter. *Non-equilibrium* or *irreversible thermodynamics* deals with processes; thus, it must be able to describe systems that can be spatially inhomogeneous and change over time. Here, the state variables no longer describe the system as a whole but become field functions depending on space and time.

## Thermodynamics of biological systems is thermodynamics of non-equilibrium open systems

### Open Systems

Biological systems must be considered as open since they exchange matter with their environment. They obtain energy in form of sun light or in form of high energy nutrients and they release “waste” of low energy. One consequence of this throughput of matter is that they cannot evolve towards equilibrium as long as they are alive. However, it is frequently observed (and used as a practical and important assumption in metabolic modeling) that they approach a state where the system’s state variables are constant despite the flow of matter. This state is typically stable and has been called steady state or, in the original version by Ludwig von Bertalanffy, “Fließgleichgewicht”^5^. Among the first to characterize these steady states thermodynamically was Ilya Prigogine^6–9^. Below we will describe important relations that are relevant for metabolic studies and cellular growth.

Here, we will first revisit the concepts of balance equations and force-flux relationships, which allow us to calculate physical properties of irreversible processes, such as reaction rates and especially entropy production and conversion. Thereafter, we will discuss the balance of entropy in open systems and their relevance for biological systems. Finally, the implications are described for a series of systems of practical importance.

### Balance Equations and Force-Flux Relationships

For open systems, we must keep track of uptake, conversion, and release of the quantities describing the states of the system such as matter, entropy or charge. Be *Y* the considered quantity, then *Y* can change by uptake from or release to the environment (indicated by index “ex” below) as well as by internal conversion (index “in”) such as chemical reactions:

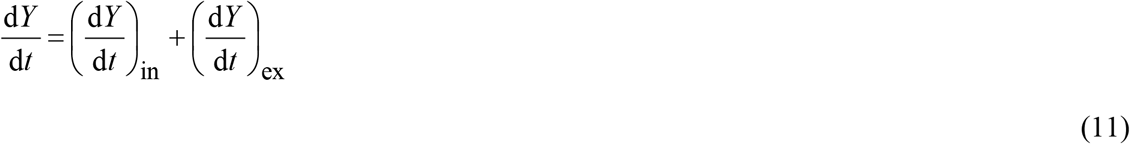

For heterogeneous systems, it is favorable to consider the density *y* instead of *Y* itself with 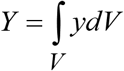 and *V* being the volume of the system. A short mathematical conversion of equation (11) results in the following local balance equation:

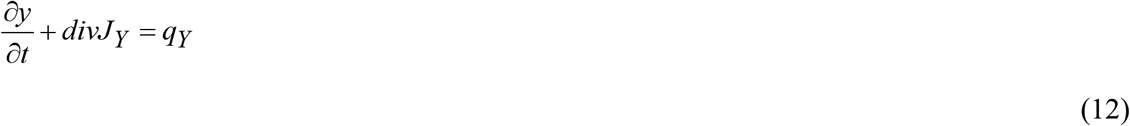

where *∂y*/*∂t* is the (local) change of *y* over time, *J*_*Y*_ is the flux density of *Y* and *q*_*Y*_ is production or decay density of *Y* at the considered point. This type of equation is known for many phenomena in technical or natural systems. If *q*_*Y*_ is zero, i.e. *Y* is neither produced nor degraded, we obtain a conservation relation. An example is the conservation of electrical charge, which can locally change through electrical current. The other way around, for a chemical compound in a closed system (test cuvette) total flux over the surface is zero 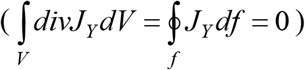 since no matter can be exchanged beyond the border (surface *f*) of the cuvette.

Irreversible Thermodynamics studies the thermodynamic fluxes *J*_*i*_. Their causes are generalized forces *X*_*k*_. For example, electrical voltage induces electrical current (Ohm’s law), a temperature gradient induces a heat flow (Fourier’s law), a concentration gradient induces a diffusion flux (Fick’s law), or a chemical affinity a conversion of matter (chemical reaction).

If several forces occur at the same time, then the quantity of every flux depends on all forces together. We don’t know in general terms how these dependencies read far away from equilibrium. However, close to equilibrium it is justified to assume (based on a Taylor series expansion) that every flux depends linearly on all forces:

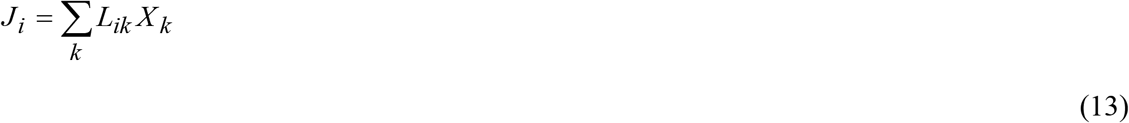

The linear coefficients *L*_*ik*_ are called Onsager coefficients^10, 11^ (after the physical chemist/theoretical physicist Lars Onsager). The diagonal elements *L*_*ik*_ with *i=k* are, for example, the coefficients of electrical and thermal conductivity, or of diffusion. They are always non-negative, i.e. the force always induces its conjugated flux. The off-diagonal elements *L*_*ik*_ with *i* ≠ *k* are the coefficients of reciprocal phenomena such as the Dufour effect (flow of heat due to concentration differences), the Sorret effect (flow of matter due to temperature gradient), the Peltier effect (flow of heat due to electrical current differences) or the Seebeck effect (electrical current due to temperature difference). In addition, we have the coefficients for electrochemical, thermoosmotic, thermochemical and other effects.

In 1931, Onsager discovered that the coefficients *L*_*ik*_ are symmetric (i.e. *L*_*ik*_ = *L*_*ki*_) close to the equilibrium, i.e. the coefficient relating e.g. the temperature gradient to diffusion is equal to the coefficient relating the concentration gradient with the heat flow.

The entropy production density within the system can be related to these forces and fluxes. It reads

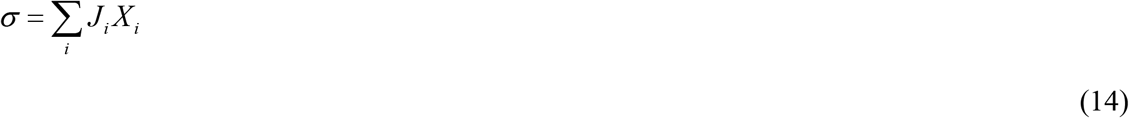

This expression is important since it covers the direction of irreversible processes in the system. The forces most interesting for biological problems are

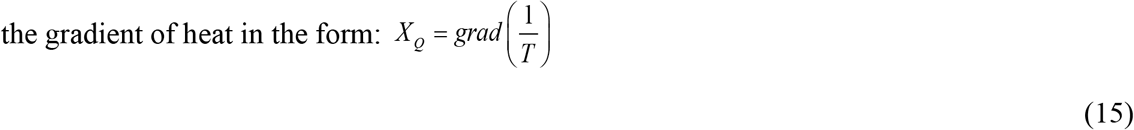

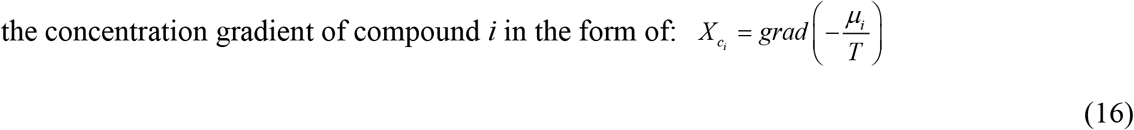

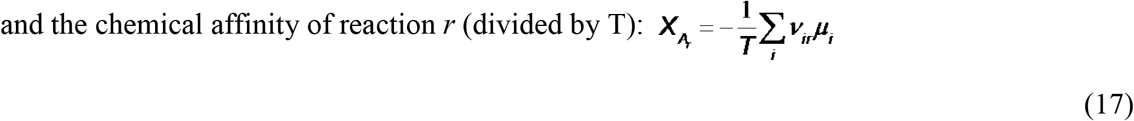

In these expressions, 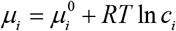 denotes the chemical potential of substance *i* with concentration *c*_*i*_. For charged molecules, the chemical potential can be replaced by the electrochemical potential 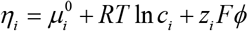, where *z*_*i*_ is the charge, *F* Faraday’s constant and *φ* the electrical potential. The *v*_*ir*_ in *X*_*A*_*r*__ are the stoichiometric coefficients of substance *i* in reaction *r*.

Combining equations (15) to (17) with equation (14), we can calculate the different fluxes in a thermodynamic system based on the actually present forces.

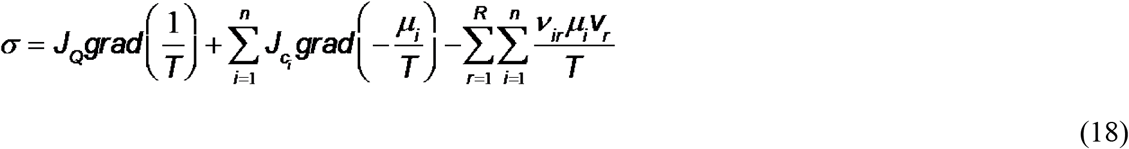

A few practical examples will be presented below. Considering in addition equation (14) allows us to determine the entropy that is produced or converted within the system during these irreversible processes.

Note that the temporal change of entropy of a system, 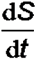, is the volume integral over *σ*. The energy dissipation due to the irreversible processes is calculated as 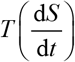. It is a measure for the energy that is no longer available for the system to perform work. Here, we can understand performing work in the sense of directed transport against a gradient instead of undirected diffusion or creation of high-energy molecules instead of their decay. Since in equilibrium the entropy assumes a maximum (and, hence, doesn’t increase further), the dissipation (as well as entropy change) vanishes in equilibrium.

### Entropy Production and Irreversibility in Open Systems – General Principles

As for any state variable, the balance equation (11) holds also for the entropy *S*, i.e.

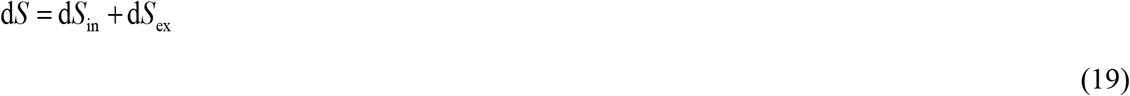

d*S*_in_ comprises the entirety of the irreversible processes. Without addition of energy from the outside of the system, d*S*_in_ is always positive or zero. It is easy to see that the unconditional increase of entropy of a system according to the second law of thermodynamics only holds in isolated macroscopic systems, since there we have d*S*_ex_= 0 and, hence, d*S*= d*S*_in_≥ 0. This principle is illustrated in Figure 1, left side.

In open systems, d*S*_ex_ can be negative due to, for example, export of heat or import of matter with low entropy or export of matter with high entropy, see Figure 1, right side. If

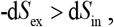

i.e. if the irreversible processes within the system do not compensate the reduction of entropy due to exchange with the environment, then also

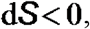

without any conflict with the second law of thermodynamics.

**Figure 1:**
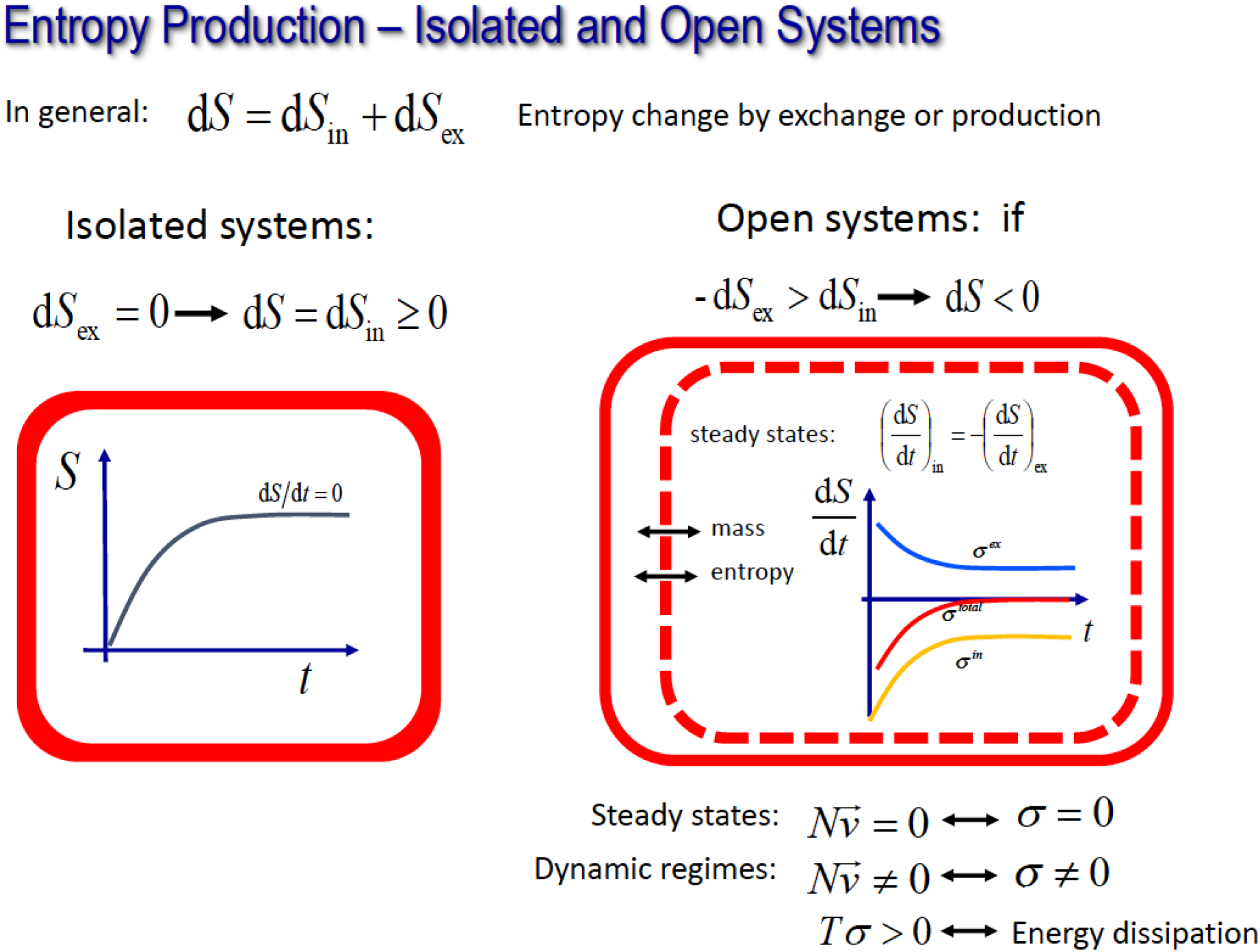
Principles of entropy production in isolated and open systems. In isolated systems, entropy is produced until they reach an equilibrium state. Then entropy is at maximum and entropy production ceases. Open systems are characterized by exchange of mass, energy, and, hence, entropy. During dynamic processes, entropy can be exchanged and, hence, in total increase or decrease. At steady state, entropy production of the full system also vanishes, ensured by a balance of entropy production in the system and entropy export.

This fact is important for biological systems since it allows for an increase in order based on the uptake of nutrients and, hence, for the development of the biological entity over time, i.e. life.

d*S*_in_ has been called a measure for the irreversibility of a physical process. It is related to the degree of irreversibility, *P*, with

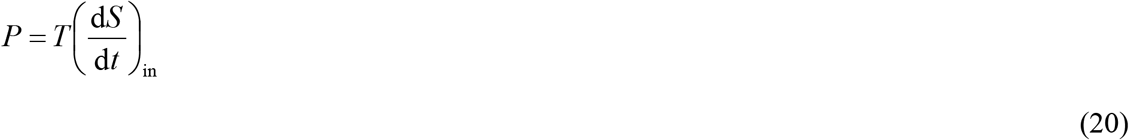

Prigogine^6^ has already found that open systems develop towards states with minimal *P*, such that d*P*/d*t* ≤ 0 and, hence,

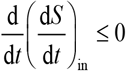

He also found that states with a minimum of *P*, which is compatible with the system’s conditions, are stable and stationary. For these states, it holds that

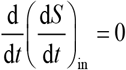

And, hence, 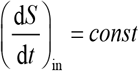.

Due to the stationarity (i.e. the constant total entropy), we find that the entropy production within the system is balanced by the export of entropy, i.e.

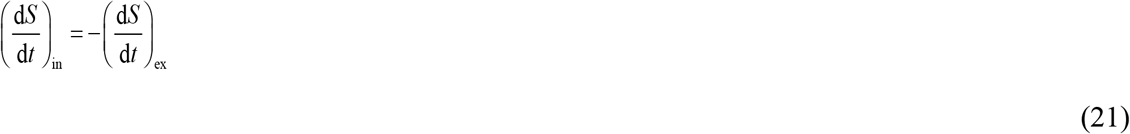

Thus, total entropy does not change and energy is not dissipated. This means that the energy provided from the outside into the system is perfectly converted into energy required for chemical reactions and build-up of high-energy molecules and cellular building blocks.

### Practical Calculation of Entropy Dynamics in Biochemical Systems

To calculate the entropy dynamics in metabolic systems, we can return to the entropy production density, introduced in equation (14) and the respective forces, equation (17), as well as the related reaction rates (or fluxes), **V**_*i*_. Neglecting heat flow and diffusion processes, the entropy production density reads

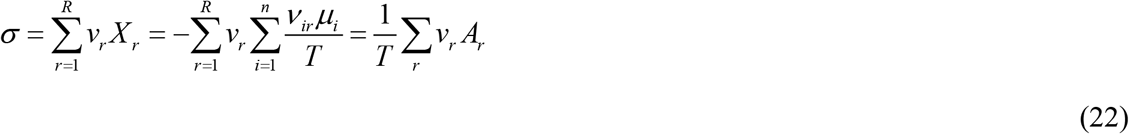

where 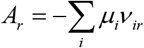 is the chemical affinity of reaction *r*. This allows for an easy calculation of the temporal changes of *σ* in any dynamical metabolic or regulatory biochemical system, as is also illustrated in Figure 2.

**Figure 2:**
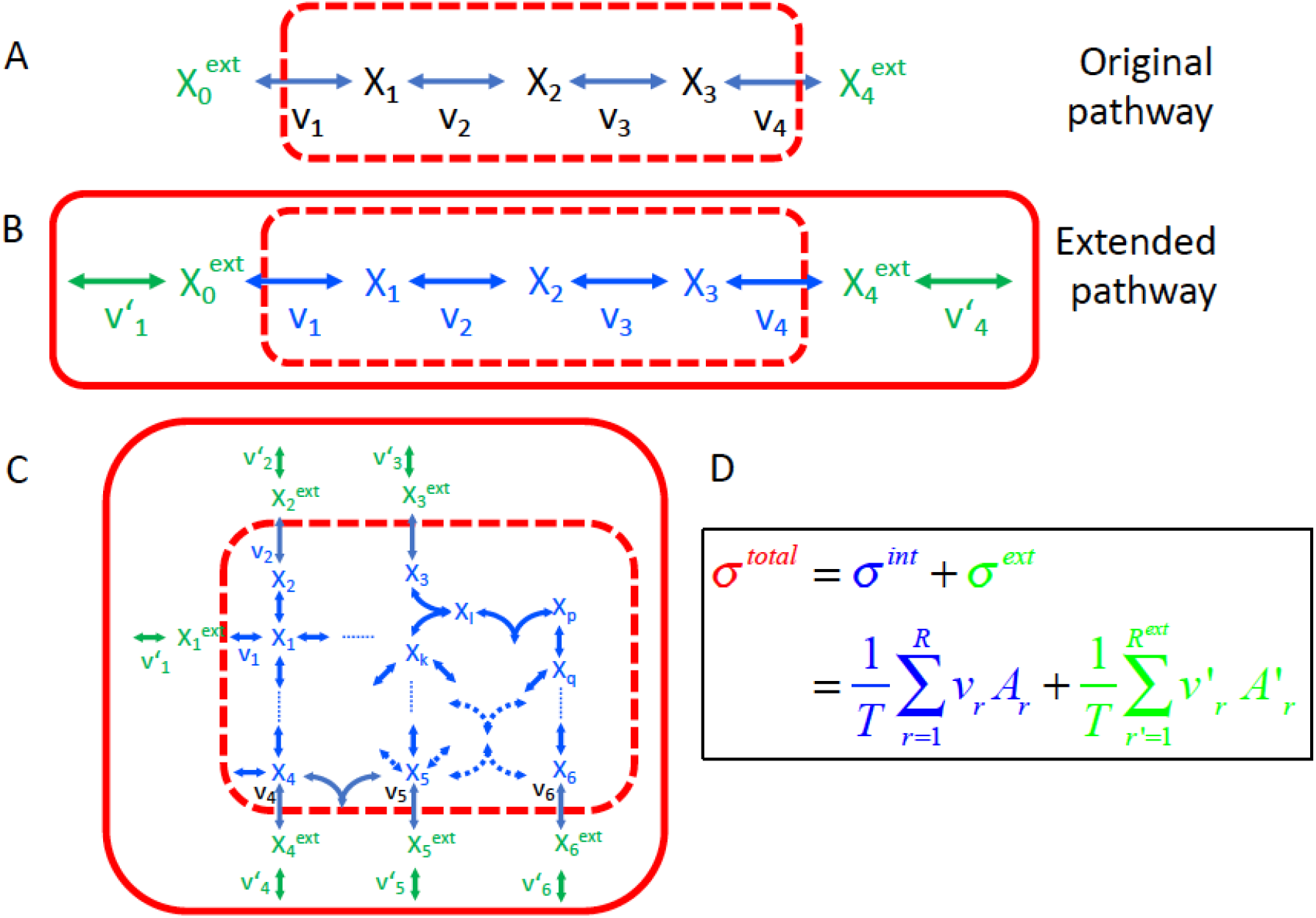
Illustrative example case for handling open systems. A) Extension of the original pathway, which is proceeding within the open system, to the extended pathway, which covers the original pathways as well as those processes ensuring constant external conditions. B) Dynamics of metabolites from an initial state towards steady state. C) Related dynamics of the reaction rates. D) Initial dynamics of entropy production terms: σ(v_i_) – entropy change rate for individual reactions, σ^intern^ – entropy change rate within the system, σ^total^ – total entropy production rate. E) same as D, only different temporal period to illustrate the behavior at steady state: while the individual reactions may have positive and negative entropy production rates, the total entropy production of extended system vanishes.

If we assume a steady state, which is of interest for systems analyzed with flux balance analysis, we also require that

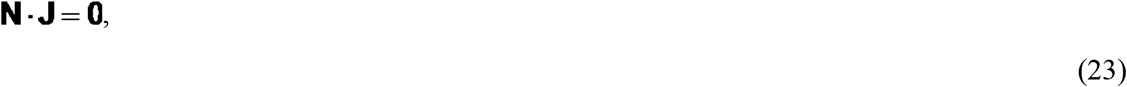

where 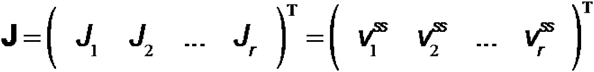 is the vector of steady fluxes and **N** = {*v*_*ir*_} is the matrix of stoichiometric coefficients. The superscript *ss* denotes steady state. Multiplying equation (23) with the row vector of chemical potentials in steady state, 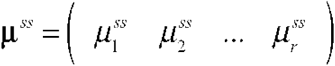 directly leads us to the fact that the entropy production density vanishes in steady states

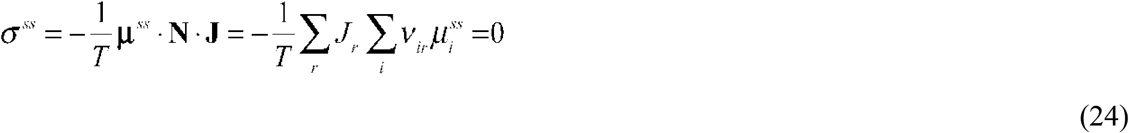

Equation (24) implies that the energy dissipation, *T*(d*S*/d*t*), of the system is also zero, i.e. the energy taken up is completely converted into the energy of, e.g., internal compounds without losses. Only away from steady state, energy of compounds taken up is partially lost as dissipated energy. Hence, steady states are not useful for biological systems to create structure, which means that growth and development have to proceed away from steady state.

### Entropy Balance in Open Systems

In open systems of biochemical reactions, we have uptake and secretion of compounds over the system’s boundaries. Some of the reactions are biochemical transport processes connecting the interior of the system with the environment. Thus, application of the equations (22) and (24) requires additional considerations, since now the separation of system and environment is no longer so obvious. In many modeling approaches, the external metabolites are considered as constant (and practically are kept constant, e.g. in a chemostat). That means that one must introduce an artificial thermodynamic replacement process that captures the entropy dynamics due to this steering process. Then, the full system to consider in the analysis is comprised of the replacement process (ensuring constant nutrients or other external metabolites), the uptake and release processes as well as the actual internal biochemical conversions. If the system is extended respectively, equations (22) and (24) keep their validity also for open system. Figure 2 illustrates the extension of the original pathway to the full (or extended) pathway as well as the resulting dynamics for metabolite concentrations, reaction rates and entropy production rates when the systems moves from initial conditions towards steady state.

### Illustrative Examples

#### 1. Chemostat and Batch culture

Let’s illustrate the dynamics of entropy production and exchange for two devices, which are of special interest for many biological investigations, the chemostat and the batch culture, both methods to culture microorganisms. The *chemostat* is a device constantly providing fresh medium and nutrients with a constant flow rate and at the same time removing waste medium and the organisms contained in the overflow. If the flow rate of the chemostat is smaller than the maximum growth rate of the cultivated microorganisms, it reaches a steady state where the growth rate of the population is equal to the wash-out rate. As a result, the number of organisms is constant. The chemostat is an open system. The *batch culture* is a closed microorganism culture system with specific initial values of nutrients, temperature and other environmental conditions. Since no further nutrients are provided and waste is not removed during the incubation, the culture can only grow until all nutrients are consumed and then growth stops.

For both cases, the dynamics of compound concentrations and entropy are exemplarily illustrated in Figure 1 (for chemostat) and Figure 2 (for batch culture). For the following consideration, we assume that the temperature is constant and no energy is provided in form of heat or work.

**Figure 3:**
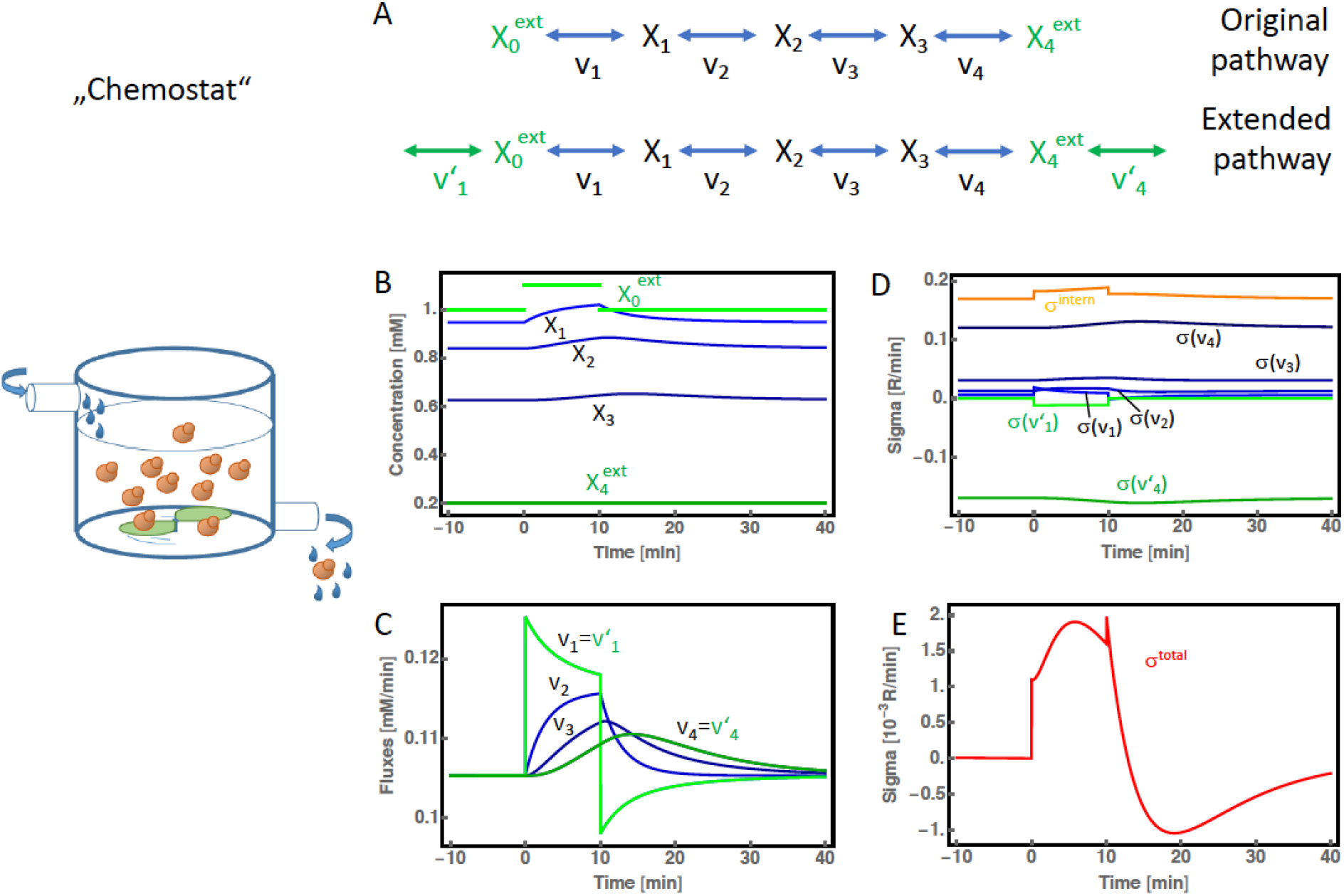
Chemostat. A) To illustrate conditions in a chemostat, we assume a simple series of metabolic reaction with X_0_ being the initial substrate and X_4_ the final product; both are kept constant by the device. To calculate the total entropy, we must extend the pathway by reactions v’_1_ and v’_4_ to account for entropy contributions due to fixation of external conditions. Since the system is normally in steady state and would not show any dynamics, we simulated a pulse of increase of external X_0_ from time *t*=10 to *t*=20. B) Time courses of metabolite concentrations. C) Fluxes through individual reactions. D) Dynamics of entropy production of individual reactions. E) Dynamics of total entropy production. Parameter values: 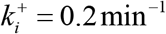, 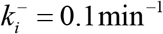, *i* = 1,..., 4

The *chemostat* practically operates in steady state. In order to simplify the consideration here, we assume only one nutrient (e.g. glucose, here denoted as X_0_), one metabolic pathway and one secretion product (here denoted as X_4_). Keeping X_0_ and X_4_ constant requires two replacement processes. In agreement with equation (24), the whole system doesn’t produce any entropy or dissipate energy as long as the chemostat is in steady state. Only if a perturbation occurs, which moves the system out of steady state, then entropy moves out of balance. In case of a glucose pulse, the individual reactions either create or consume entropy, depending on the amount signs of reaction rates and the chemical potentials of educts and products over time. In total, the system produces entropy until it returns to steady state where entropy production again vanishes. When the additional glucose supply is turned off, the overall even assumes a negative sign, the system exports more entropy than it imports.

In the *batch* culture, we can distinguish three main phases: In the first phase or *lag phase*, we have an excess supply of nutrients, but only few cells to make use of them. In the second phase or phase of *exponential growth*, nutrient uptake and growth are balanced. Since nutrients are not yet limiting, the culture reaches a quasi-steady state. In the third phase or *stationary phase*, nutrients become depleted and growth will finally stop. For a while, growing cells will feed from dying cells. Eventually, the whole population will die out and become degraded. The culture develops towards thermodynamic equilibrium.

For the batch culture it is important to precisely state what we consider as system. Two possible choices are (i) the whole batch as closed system or (ii) the cell population as open system with the batch as environment. For the first case, we have d*S*_ex_ = 0 and d*S* = d*S*_in_ ≥ 0 (until equilibrium is reached).

For the second case, we have uptake of Gibbs enthalpy and export of entropy in the first phase, steady state conditions as described by equations (22) for the second phase, and an increase of entropy until its maximum after the nutrients have been depleted.

**Figure 4:**
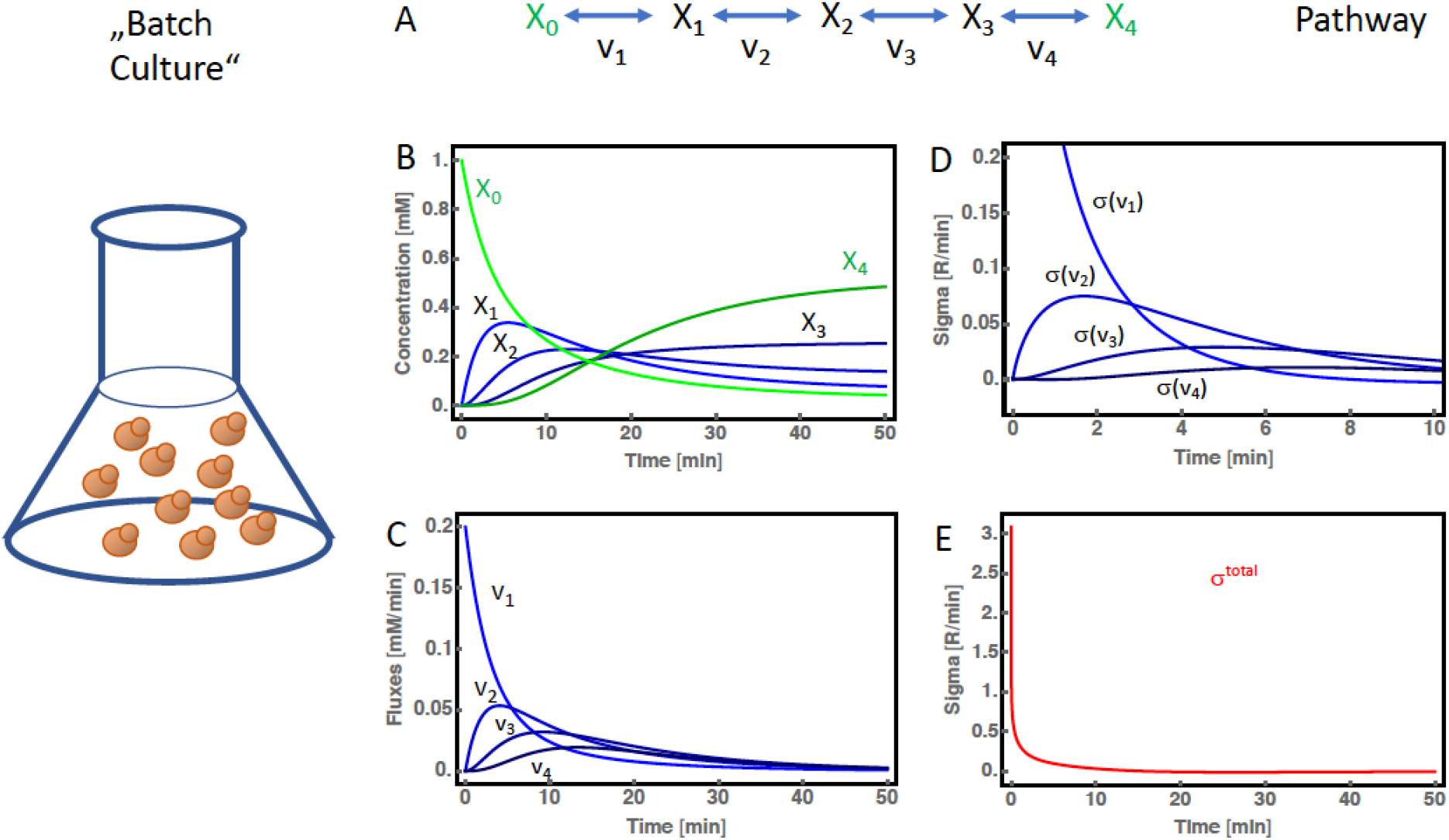
Batch culture. A) In a batch culture, the initial substrate X_0_ will be consumed and fully converted by the chemical reactions into the final product (or waste) X_4_. No extensions of the systems are required to calculate the total entropy. B) Time courses of metabolite concentrations. C) Fluxes through individual reactions. D) Dynamics of entropy production of individual reactions. E) Dynamics of total entropy production. Parameter values: 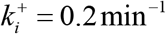, 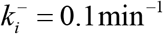, *i* = 1,..., 4

### 2. Metabolic Model for healthy cells and cells with cancer mutations

In a recent study, the impact of mutations in signaling pathways on the behavior of central carbon metabolism has been studied^12^. The mutations are characteristic for many cancers. The metabolic model, which is fitted to data for healthy and mutated cells, is used here for further analysis (Figure 5). Cells have been grown under conditions of practically constant glucose concentration. Several compounds are secreted, such as lactate, glucose-6-phosphate and serine. Again, we have to assume replacement reactions that keep both the external glucose constant and the secreted compounds in the environment on a zero level.

**Figure 5.**
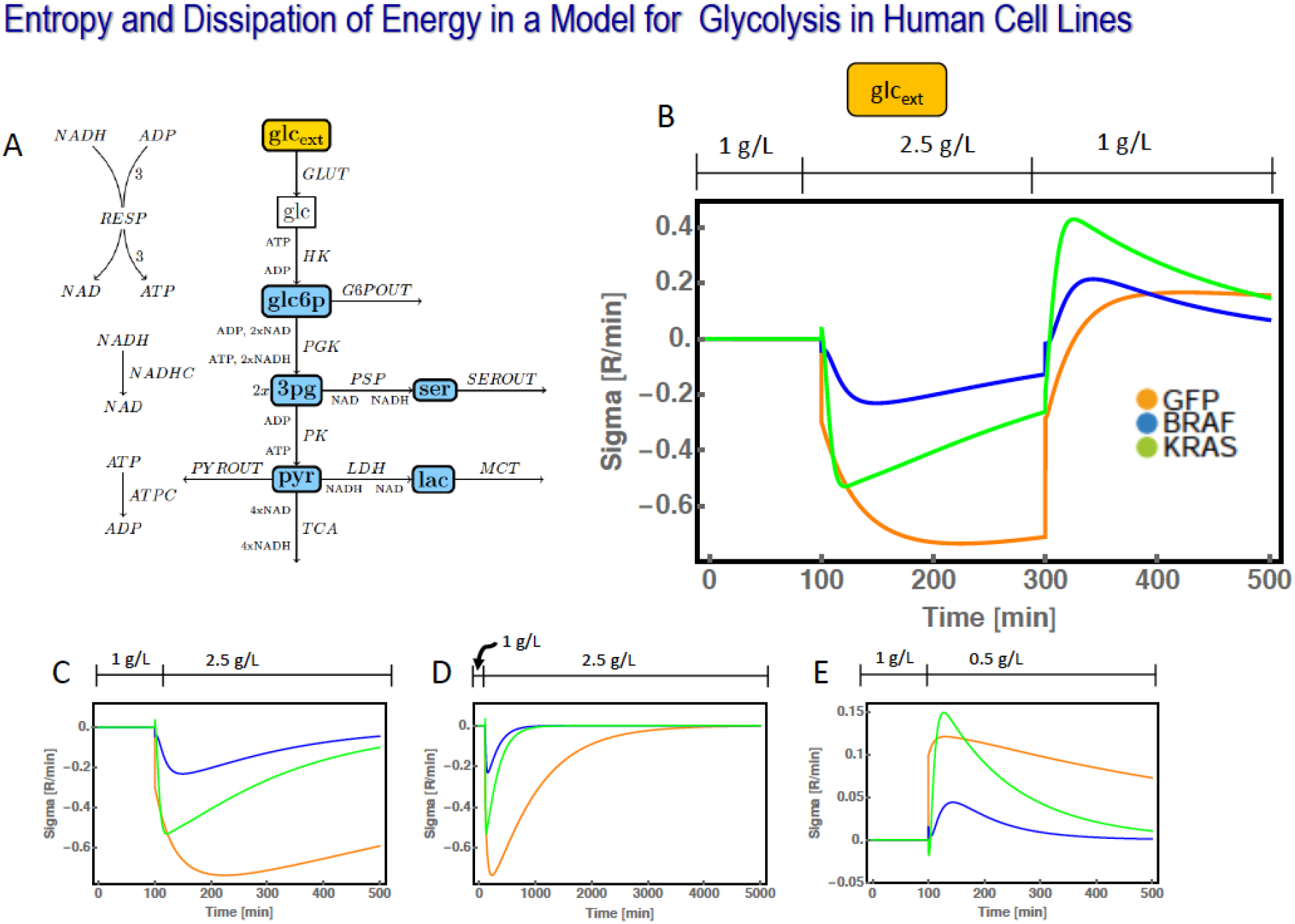
Entropy dynamics in a model for glycolysis in different human cell lines (GFP – wild type expressing GFP, BRAF – Braf mutant, KRAS – Kras mutant). A) Wiring scheme of the network. B) Comparative dynamics of entropy production for the three strains during upshipt and downshift of external glucose. C) Dynamics of entropy production upon glucose upshift only. D) Longterm dynamics after glucose upshift. E) Dynamics of entropy production upon glucose downshift. Dynamics of metabolite concentrations, reaction rates and entropy production rates related to individual reactions are shown in Supplementary Information.

Here, like in the above considered cases, entropy export and production are balanced as long as the system is in steady state. When the system is perturbed, e.g. by increasing the external glucose concentration from 1 g/L to 2.5. g/L (changes done in the experiments), then all reactions show either positive or negative contributions to *σ*. It is interesting to note that increase of *σ* is by far more pronounced in wild type compared to the cancer cells. Though is requires further investigation, it might be a hint towards an organization of metabolism in cancer cells that allows for more effective use of the provided energy and prevents the higher level of energy dissipation in healthy cells.

When the glucose level is decreased, *σ* assumes negative values, indicating that more entropy is exported than produced by the cells.

### 3. Dependency on kinetics illustrated for a simple metabolic pathway

A simple model of a metabolic pathway can also be used to illustrate a few more important dependencies of entropy dynamics on the kinetics of such a pathway. First, we can ask for the impact of the speed of individual reactions on entropy production in case of perturbations. We explicitly consider a case where the overall drop in Gibbs energy (or equally, the overall equilibrium constant) is the same, but the realization in terms of kinetic constants of forward and backward reactions differs. Here, we have to distinguish between instantaneous or short-time behavior and infinite times. On short time scales, fast reactions (with high forward and backward kinetic constants) show higher entropy production than slow reactions. If we integrate the entropy production from the time point of perturbation until infinity, then fast and slow reactions produce in total the same amount of entropy after identical perturbations. This is illustrated in Figure 6.

**Figure 6:**
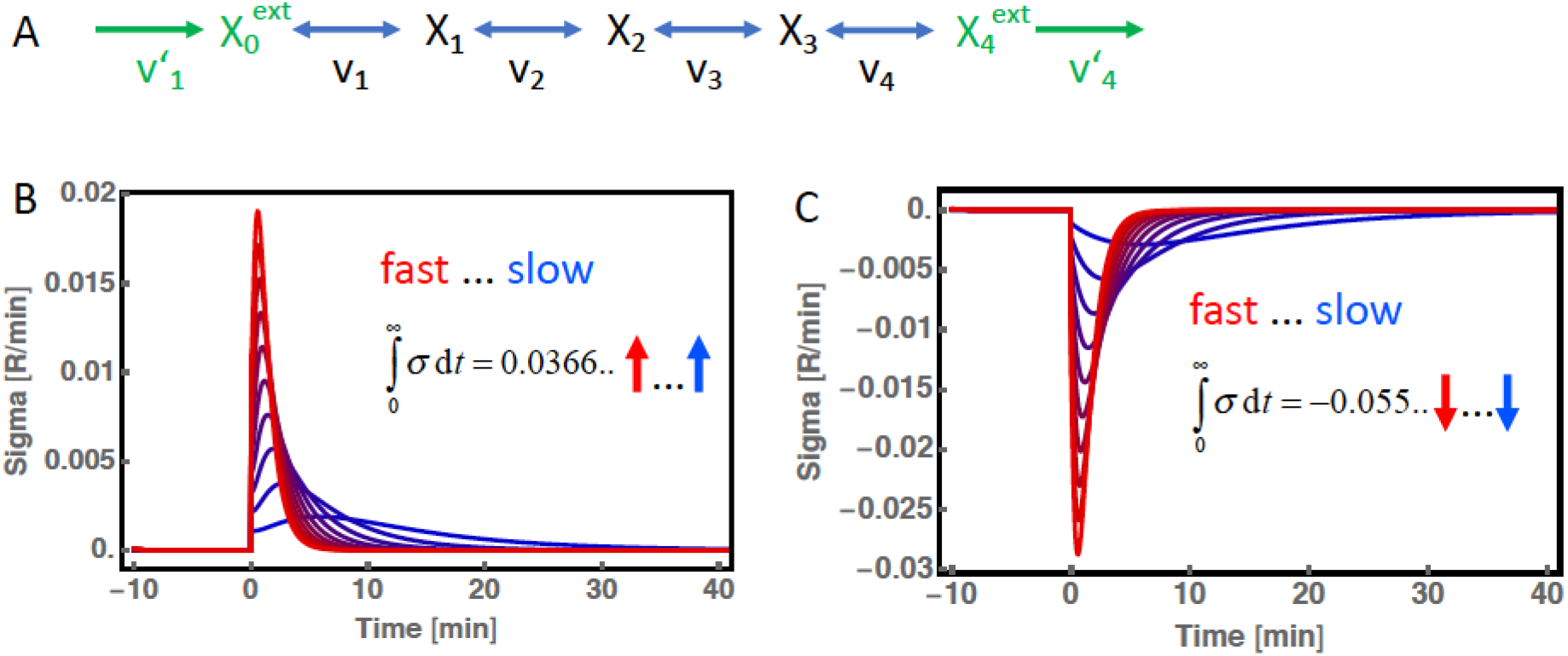
Faster reactions produce more entropy than slow reactions on short time scales, but the same total entropy on long time scales. A) Network, B) Total entropy dynamics for fast (red curves) and slow reactions. The perturbation is an increase in 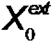 and the system returns to a steady state. While the entropy production rate differs depending on the speed of the reaction, the total entropy production until return to steady state is always the same (about 0.0366 R) C) Total entropy dynamics upon decrease in 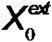. Parameter values 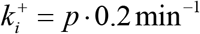, 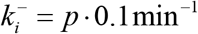, *i* = 1,..., 4, *p* = 1...10.

### Entropy production in systems with limit cycle oscillations

Systems showing dynamic behavior are not in steady state and exhibit dynamics in entropy production. That also holds for systems with oscillations. However, when the system has reached the perfect limit cycle, then entropy production and entropy degradation over one cycle are balanced such that the integral, i.e. the total production, for one period vanishes. Figure 7 illustrates that behavior for the example of the Higgins-Sel’kov oscillator. Higgins and Sel’kov had suggested this model to explain oscillations observed in the glycolysis of yeast cells as consequence of a positive feedback from Fructose-1,6-bisphosphate (here presented by compound Y) on its own production by the enzyme phosphofructokinase. The rates are

**Figure 7:**
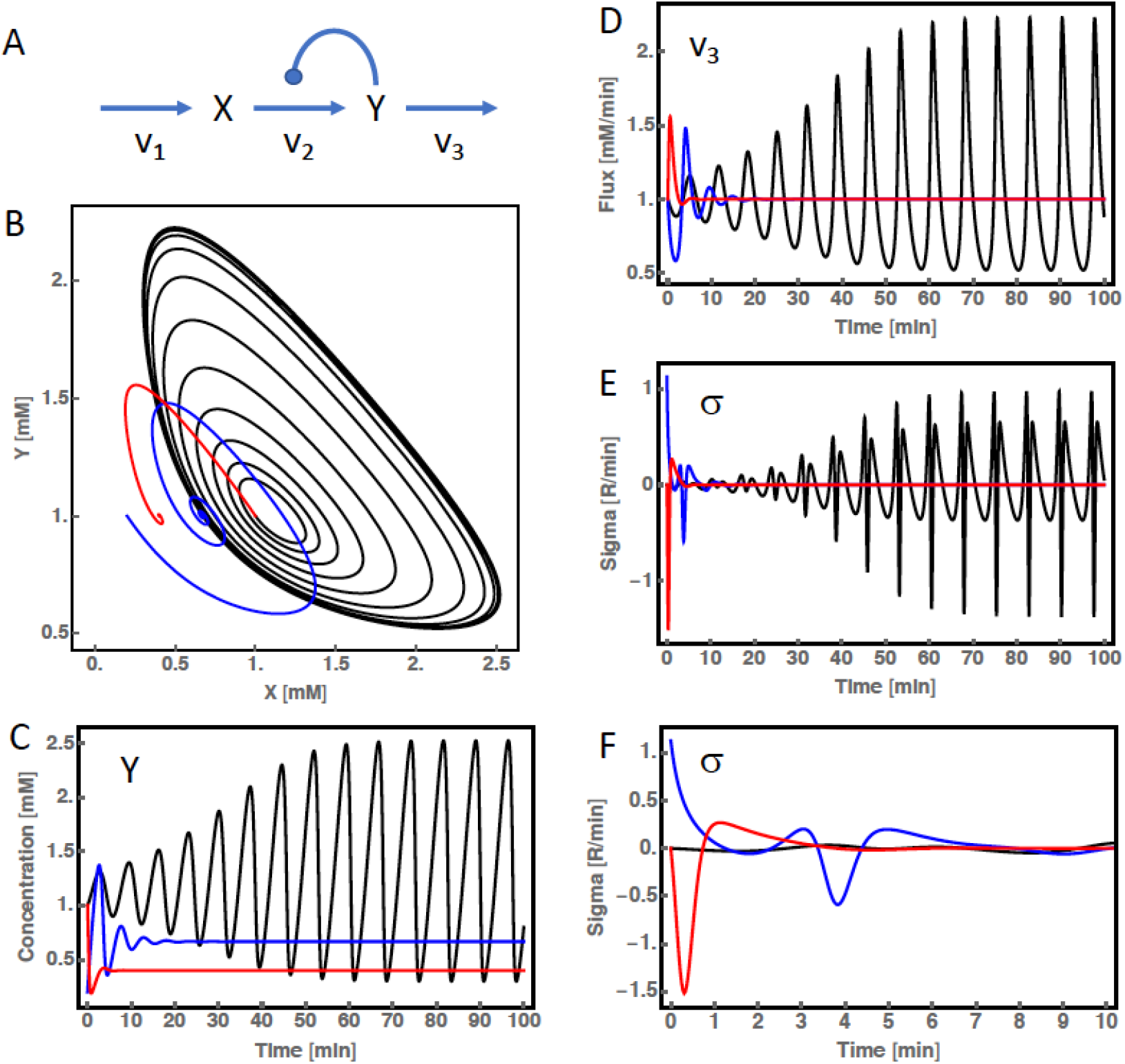
Oscillations and entropy production. A) The classical Higgins-Sel’kov oscillator was used as model system. B) Phase plot of the concentration dynamics for different parameters and initial conditions (red – node, blue – damped oscillations, black – limit cycle oscillations). C) Time course of component Y, D) Time course of rate v_3_. E+F) Entropy production for the different cases, in different temporal resolution. Parameter values: *v*_1_ = 1mM ⋅ min^−1^, *k*_2_= 1min^−1^, black: *k*_3_ = 0.9 min^−1^, blue: *k*_3_= 1.5 min^−1^, red: *k*_3_ = 2.5 min^−1^

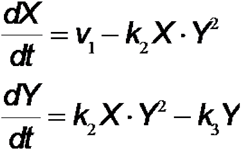

Depending on the values of the fixed influx rate v1 as well as of the parameters k2 and k3, the system can evolve towards a steady state in form of a node or in form of a focus (damped oscillations) or it can exhibit limit cycle oscillations.

## Conclusions from the Examples

The entropy production rate as introduced in Equation (22) together with considerations about boundary conditions and entropy provided or depleted by processes keeping boundary conditions constant, allows for calculation of the energy expenditures in metabolic processes. As demonstrated, the dynamics of entropy production depend on the network structure, the kinetic details and on the initial conditions of the system. If the system is allowed to evolve towards a steady state, entropy production vanishes. As a rule of thumb, the entropy production rate (in absolute terms) is higher for faster processes. However, overall entropy production does not depend on the speed of individual reactions, but on the overall drop in Gibbs free energy of a pathway or network. As a consequence, for oscillating systems exhibiting limit cycle oscillations, the overall entropy production per cycle is zero. However, during the transient, the production rate can be positive, negative or show damped oscillations.

Equation (22) can also be applied to subnetworks of a large, e.g. genome-scale, metabolic network model to assess how much entropy individual subnetworks produce or consume. This would give hints to optimization potential, e.g. in biosynthetic production processes, or to areas interesting for regulation in healthy and diseased cells. However, to this end, one first needs a metabolic model with sufficiently accurately determined kinetics.

### Beyond Culture Conditions

Concepts of non-equilibrium thermodynamics and the relation between forces and fluxes have been fruitfully applied to a series of biological problems. For example, it has been analyzed whether the order of ATP producing and consuming reactions in glycolysis makes a difference for the glycolytic flux and the energy yield of glycolysis^13^. It could be shown that it is advantageous in terms of maximizing the glycolytic flux to have ATP consuming reactions at the beginning and ATP producing reactions at the end of the pathway, respectively. Ion transport over biological membranes is typically described with force-flux relationship (e.g. ^14, 15^) and the coupling of the transport of different ions with consideration of membrane potential has recently been tackled by such an approach^16^.

For the very active field of modeling metabolic networks it is of outmost importance to take thermodynamic constraints into account. This is by now a commonality in flux balance analysis ^4, 17, 18^. A series of attempts have been made to estimate thermodynamically consistent parameters in enzyme kinetic models such as parameter balancing^19^ or the assignment of Δ_*r*_*G* to enzyme kinetics ^20–22^. However, there have also been some misconceptions about how to apply thermodynamics to metabolic networks. For example, it had been suggested that minimizing energy dissipation (in systems analyzed under the assumption of steady state) could be a governing principle for such networks [Ref Heinemann]; as demonstrated above that is physically incorrect since energy dissipation vanishes in steady state.

Overall, we must be aware of the fact that flux balance and the related constraint concepts only hold in steady state. Away from steady state, i.e. during active growth and development, during response to stress, during oscillatory regimes and essentially in most situations characteristic for live, biological systems have to face energy dissipation and dynamics in entropy production in order to create new structure.

## Acknowledgments

This work has been funded by the Deutsche Forschungsgemeinschaft (DFG, TRR 175 “Greenhub”). We thank Dr. Martin Seeger for critical reading.

